# CellExplorer: a graphical user interface and a standardized pipeline for visualizing and characterizing single neurons

**DOI:** 10.1101/2020.05.07.083436

**Authors:** Peter C. Petersen, Joshua H. Siegle, Nicholas A. Steinmetz, Sara Mahallati, György Buzsáki

## Abstract

The large diversity of neuron types of the brain, characterized by a unique set of electrophysiological characteristics, provides the means by which cortical circuits perform complex operations. To quantify, compare, and visualize the functional features of single neurons, we have developed the open-source framework, CellExplorer. It consists of three components: a processing module that calculates standardized physiological metrics, performs neuron type classification and detects putative monosynaptic connections, a flexible data structure, and a powerful graphical interface. The graphical interface makes it possible to explore any combination of pre-computed features at the speed of a mouse click. The CellExplorer framework allows users to process and relate their data to a growing collection of “ground truth” neurons from different genetic lines, as well as to tens of thousands of single neurons collected across our labs. We believe CellExplorer will accelerate the linking of physiological properties of single neurons in the intact brain to genetically identified types.

## INTRODUCTION

Discovering novel mechanisms in brain circuits requires high-resolution monitoring of the constituent neurons and understanding the nature of their interactions. Identification and manipulation of different neuron types in the behaving animal is a prerequisite for deciphering their role in circuit dynamics and behavior. Yet, currently, a large gap exists between neuron classification schemes based on molecular and physiological methods (Gouwens et al., 2019; Jia et al., 2019; Kepecs and Fishell, 2014; Klausberger and Somogyi, 2008; McBain and Fisahn, 2001; Roux and Buzsáki, 2015; Rudy et al., 2011).

Large-scale extracellular electrophysiology aims to establish the relationship between neuronal firing and behavioral or cognitive variables in order to provide insights about the computational role of neurons and neuronal assemblies (Barlow, 1972; Buzsáki, 2004; Steinmetz et al., 2019). However, exploiting the power of correlations between neuronal firing and behavioral variables requires multi-level characterization of single neurons and their interactions. Simultaneous recordings from large numbers of neurons, preferably identified by optogenetic and other methods, make it possible to build an extensive list of neuron feature and their assigned ‘cell type’ properties (Fig. 1). These properties can be described at multiple levels of complexity. The first level is a description of the biophysical and physiological characteristics of single neurons. This level includes waveform features (Fig. 1B), their position relative to the recording sites and other units (Csicsvari et al., 2003), interspike interval statistics, and autocorrelograms (Fig. 1C). These first-level features can be used for the first-order separation of single neurons into putative major classes, typically excitatory and inhibitory cells (Fig. 1D). The second level relates single neurons to other neurons and includes cross-correlations and putative monosynaptic connections derived from spike transmission probabilities (Fig. 1E), relationship to multiple oscillatory and irregular local field potentials (LFP; e.g., rhythmic patterns, sharp-wave bursts, up-down transitions in cortex). The third level metrics of single-unit activity include the relationship between its firing patterns and brain states (e.g. non-REM, REM, awake; Fig. 1H) and overt behavioral correlates (Fig. 1I), including spontaneous motor patterns and autonomic parameters (McGinley et al., 2015; Stringer et al., 2019). In turn, these physiological properties can be related to genetically identified neuron classes with optogenetics methods (Boyden et al., 2005; Klausberger and Somogyi, 2008; Rudy et al., 2011; Buzsáki et al., 2015; Roux and Buzsáki, 2015). Antidromic and unit-LFP coupling techniques provide further assignment of single neurons to cortical regions, layers, and target projections (Bishop et al., 1962; Zhang et al., 2013; Ciocchi et al., 2015; Senzai et al., 2019).

**Figure 1:**
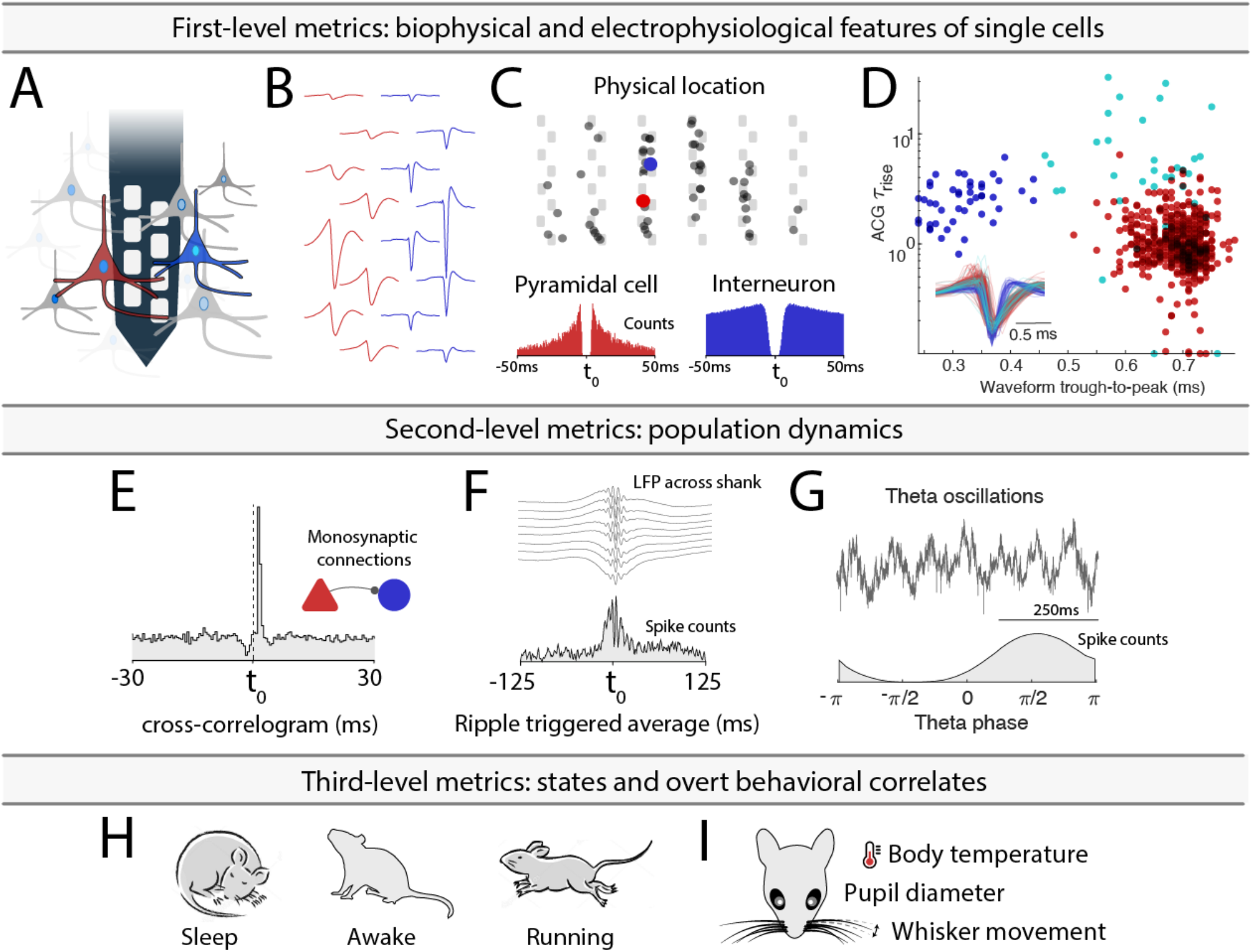
Multifaceted single neuron characterization. **A.** Using high-density silicon probes or multiple tetrodes (shown is a single shank with 8 recording sites), dozens to hundreds of neurons can be recorded simultaneously. **B**, Spikes of putative single neurons are extracted from the recorded traces and assigned to individual neurons through spike sorting algorithms, their relative position determined through trilateration (representation shows neurons projected on a silicon probe with 6 shanks) and autocorrelograms (ACGs) are used to characterize the neurons (bursting pyramidal cell in red; fast spiking interneuron in blue). **D**. Neuron-type classification based on first-order physiological parameters, using the spike waveform width (trough-to-peak) and the temporal scale of the ACGs (**τ**_rise_). Optogenetic and other direct identification methods can ground units to neuron types. **E**. Interactions between neurons are characterized by their cross-correlograms and monosynaptic connections (determined via spike transmission probabilities). **F**. Relating spikes to LFP patterns, e.g. to sharp wave ripples. **G.** Relationship to oscillations, e.g. theta oscillations (7-12Hz) with a phase histogram below from a CA1 neuron. **H-I**. Spike pattern correlations with brain states and overt behaviors.

These three levels provide universal features of neuronal activity common to all experimental paradigms and, therefore, are communicable across different experiments and laboratories. In turn, these universal features can be contrasted and compared with experimenter-provided manipulations and higher-level correlates. Because these latter variables are often paradigm-specific and differ across laboratories, the three-level analysis can guard against mistakenly assigning cognitive roles of neuronal spiking that may be explained by measurable overt correlates. Yet, even if all of the above information is available separately, factoring out critical variables and their combinations is possible only when the multitudes of single neuron characteristics can be compared flexibly.

Whether testing a specific hypothesis or mining the ever-growing number of publicly available datasets, the process can be advanced by fast and user-friendly visualization methods. To this end, we developed the open-source MATLAB based framework, *CellExplorer*, to characterize and classify single-cell, i.e. neuron, features from multi-site extracellular recordings. It consists of a pipeline for extracting and calculating physiological features, a flexible data format, and a powerful graphical interface that allows for fast manual curation and feature exploration.

## RESULTS

The CellExplorer architecture and operation consist of three main parts: a processing module for feature extraction, a graphical interface for manual curation and exploration, and a standardized yet flexible data structure (Fig 2). A step-by-step tutorial is available in the supplementary section, and more tutorials are available online (Suppl. Video 1). Flow charts are available in Suppl. Figure 2. The first step in running the pipeline is defining the data input.

**Figure 2.**
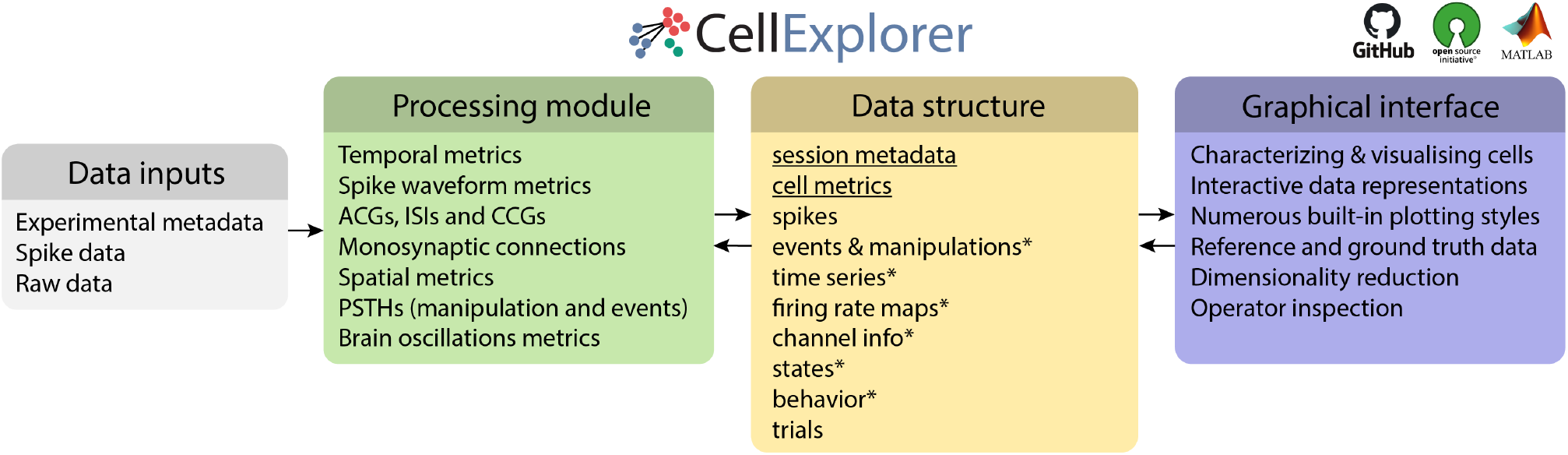
Three-component framework. A single extensive processing module (green); Standardized yet flexible data structure (yellow); and a graphical interface (purple). Data inputs are compatible with most existing spike sorting algorithms (grey). The data structure joins the Processing module with the Graphical interface (* signifies data containers). CellExplorer is open-source, built in MATLAB, and available on GitHub.

### Data Input

Before running the pipeline, relevant metadata describing the spike format, raw data, and experimental metadata must be defined (Fig. 2). All experimental metadata (session-level) are handled in a single MATLAB structure, with an optional GUI for manual entry. The platform supports several spike sorting data formats, including Neurosuite, Phy, KiloSort, SpyKING Circus, Wave_Clus, MClust, AllenSDK, NWB, ALF, MountainSort, IronClust, (Chung et al., 2017; Hazan et al., 2006; Pachitariu et al., 2016; Quiroga et al., 2004; Schmitzer-Torbert et al., 2005; Yger et al., 2018). The raw data (wide-band) is critical for comparing derived metrics across laboratories since preprocessing pipelines vary widely and depend on equipment type and filter settings.

### Processing Module

From the input data, the processing module will generate cell metrics corresponding to the three-level description of neuronal firing and their relationship to experiment-specific behaviors (Figure 2; Suppl. Table 2 contains an incomplete list of metrics for illustration; full list available at CellExplorer.org). The processing module is comprised of a single MATLAB script, *ProcessCellMetrics.m,* which computes metrics using a modular structure. The first-level description provides temporal features, waveform features (filtered and wideband), interspike interval statistics (ISIs), and autocorrelograms (ACGs). Next, the unit parameters are used for the initial classification of single neurons into broad default classes: putative pyramidal cells, narrow waveform interneurons, and wide waveform interneurons. In experiments with silicon probes, the physical position relative to recording sites is also determined using trilateration (Petersen and Berg, 2016; Csicsvari et al., 2003).

The second level of description relates single neuron spikes to the activity of other neurons and population patterns. These metrics include spike cross-correlograms (CCGs), quantitative identification of putative monosynaptic connections, phase relationships to various local field potential (LFP) patterns, and to unit population patterns. Monosynaptic connections, in turn, can be used to identify putative excitatory and inhibitory neurons and use this information to refine the primary unit classification (Fig. 1E; Suppl Fig. 3H; Barthó et al., 2004; English et al., 2017). All parameters can be customized according to the needs of each experimental paradigm (Suppl Table 2; CellExplorer.org/datastructure/standard-cell-metrics).

The third-level metrics are used to assess the relationship between firing patterns of neurons and overt behaviors, including immobility, locomotion, and running speed. Level 1-3 metrics can further be supported by optogenetic methods, which can bind physiological parameters to genetically identified neuron groups (Boyden et al., 2005; Buzsáki et al., 2015; Roux and Buzsáki, 2015). Because these 3-level metrics of single unit features are universal, they can be readily compared with similar analyses across laboratories, independent of paradigm-specific features. Towards these goals, the processing module automatically generates all cell metrics in a standardized fashion.

Features related to any behavioral paradigms, can also be computed, including manipulations (PSTHs), behavioral tracking (spatial firing rate maps) and task related trial-wise response curves (e.g., response to a sensory cue).

### Data structure

The data structure of CellExplorer (summarized in Fig. 2 and supplementary Fig. 1) is organized in data containers and MATLAB *structured arrays* (“structs”), which functionally separate different data content, making them both easily interpretable (human-readable) and machine-readable. The format is derived from the Buzcode (a MATLAB based data format for electrophysiological recordings and toolset developed communally in the Buzsaki lab; github.com/buzsakilab/buzcode), Neurosuite (neurosuite.sourceforge.net), and Freely Moving Animal (FMA) Toolboxes (fmatoolbox.sourceforge.net).

The two most important structures are the **session** metadata struct and **cell_metrics** struct.

#### The session metadata struct

Contains all session-level experimental metadata (Suppl fig 1). A session is defined as a set of data typically recorded with the same day in the same subject and is also commonly referred to as a single dataset. The **metadata struct** has a modular structure that makes it flexible and expandable, intuitive and interpretable, and it offers a single structure preventing scattering of metadata. A GUI (*gui_session.m*) allows for intuitive manual metadata entry, and a template script (*sessionTemplate.m*) can assist in importing existing experimental metadata. See https://cellexplorer.org/datastructure/data-structure-and-format/ for more info.

#### The cell_metrics struct

Modular structure containing all cell metrics calculated in the processing module. It consists of three types of data-fields for handling the diverse types of data: *numeric double*, *character-cells,* and *structs*. Single value metrics are stored in double and character cells for respective numeric and character metrics. Time series (e.g., waveforms), group data (e.g. synaptic connections), and session parameters are stored in predefined struct modules. This structure makes the content machine-readable, including user-defined metrics, and provides expandability and flexibility, while maintaining compatibility with the graphical interface. The single struct allows for processing multiple sessions together in the graphical interface (batch processing) and is convenient for sharing with collaborators and the broader scientific community in publications (see Supplementary Section and Supplementary table 2 for a detailed description and https://cellexplorer.org/datastructure/standard-cell-metrics/

### Graphical Interface

The most important component of the framework is the user-friendly graphical interface (Fig. 3), which allows for characterization and exploration of all single unit metrics through a rich set of high-quality built-in plots, interaction modes, neuron grouping, cross-level pointers, and filters. User-defined numbers of plots can be selected at any time and replaced on the screen instantaneously. In the typical layout, the top row displays population-level representations, and the bottom row displays single cell features. Selecting individual neurons from any plot automatically updates their features in the remaining interface. For easy navigation and selection, the left mouse click links to the selected neuron and the right mouse click selects the neuron(s) from any of the plots. These selected groups can be displayed alone or highlighted and superimposed against all data in the same session, multiple sessions, or the entire database. Clusters of neurons of interest can be selected by drawing polygons with the mouse cursor, and the features of the selected groups can be shown separately. Multiple group selections are also possible for both visualization and statistical comparison. Flexibility is assisted by self-explanatory side panels. The left side panel contains a custom plot panel, color group panel, display settings panel, and legend. The right panel contains single cell actions including a navigation panel, cell assignment panel, tags, and a table with metrics. The left panel also includes a text field for custom text filters and there is a message log below the main plots. The group plotting options include 2D-representation, 3D-representation, raincloud plot, t-SNE, and double histogram. Axis scaling can be either linear or logarithmic (Fig. 3E).

**Figure 3:**
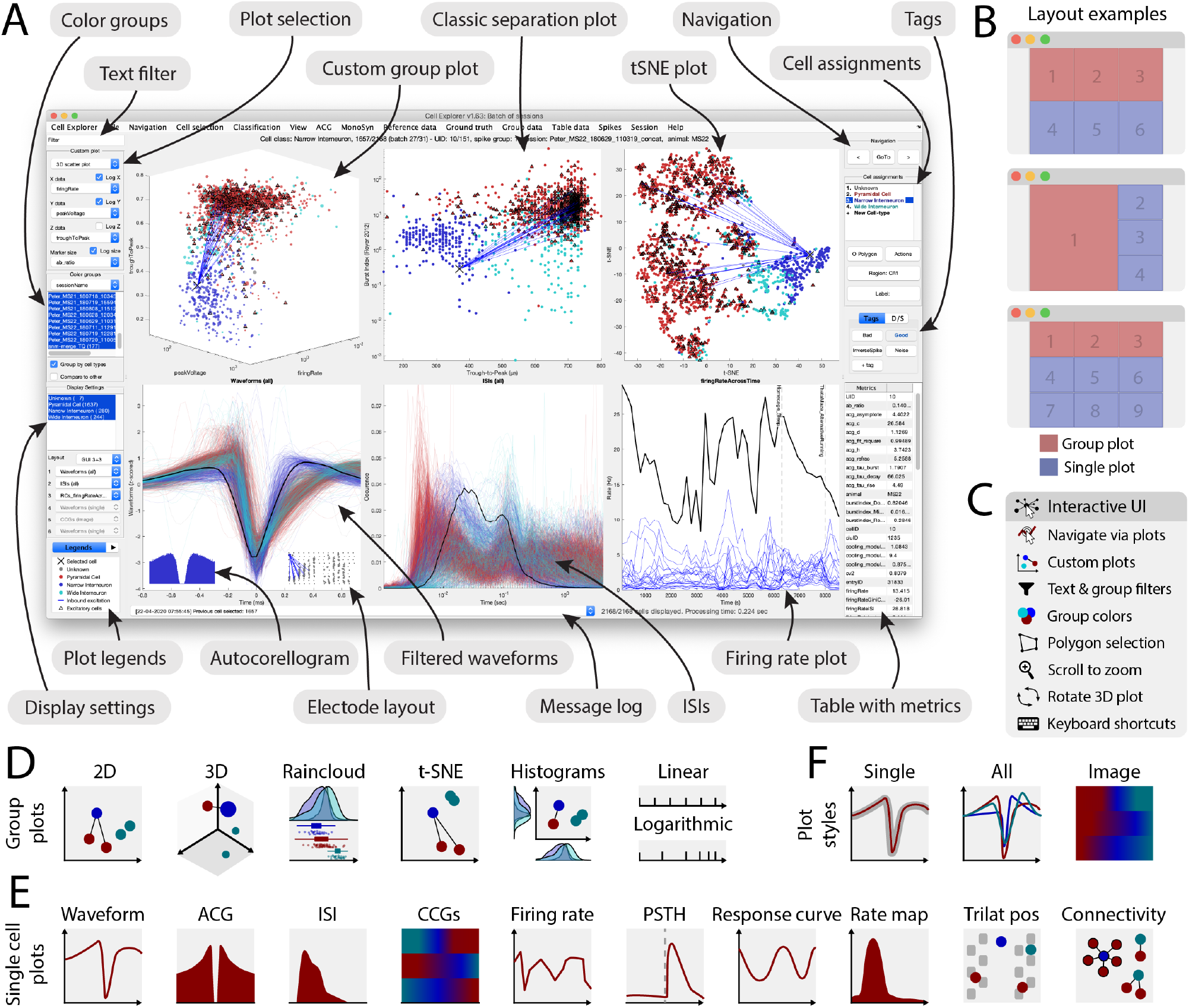
Graphical interface. **A.** The interface consists of 4 to 9 main plots, where the top row is dedicated to population-level representations of the neurons. Other plots are selectable and customizable for individual neuron (e.g., single waveforms, ACGs, ISIs, CCGs, PSTHs, response curves, and firing rate maps). The surrounding interface consists of panels placed on either side of the graphs. The left side displays settings and population settings, including a custom plot panel, color group panel, display settings panel, and legends. The right side-panel displays single-cell dimensions, including a navigation panel, neuron assignment panel, tags, and a table with metrics. In addition, there are text fields for a custom text filter and a message log. **B.** Layout examples highlighting three configurations with 1-3 group plots and 3-6 single neuron plots. **C.** The interface has many interactive elements, including navigation and selection from plots (left mouse click links to selected cell and right mouse click selects the neuron from all the plots), visualization of monosynaptic connections, various data plotting styles (more than 30+ unique plots built-in), supports custom plots; plotting filters can be applied by text or selection, keyboard shortcuts, zooming any plot by mouse-scrolling and polygon selection of neurons **D.** Group plotting options: 2D, 3D, raincloud plot, t-SNE, and double histogram. Each dimension can be plotted on linear or logarithmic axes. **E.** Single cell plot options: waveform, ACG, ISI, firing rate across time, PSTH, response curve, firing rate maps, neuron position triangulation relative to recording sites, and monosynaptic connectivity graph. **F.** Most single cell plots have three representations: individual single cell representation, single cell together with the entire population with absolute amplitude and a normalized image representation (colormap).

Examples of the flexible operation of the graphical interface module are illustrated in Fig. 4 and described in more detail in Supplementary video 1. Here we begin with motifs of monosynaptically connected clusters of neurons from the hippocampal CA1 area, as provided by the Processing Module (Fig. 4A). An example sub-network of connected neurons is highlighted in panel B with a selected single neuron (arrow) to be characterized. Selected levels one, two, and three metrics of the neuron are displayed in panels C to G, respectively. In several panels, the metrics of the selected neuron are shown against other neurons from the same dataset. Left mouse clicking on any neuron will update all the panels, allowing quick screening and qualitative evaluation of multiple features of each chosen neuron. Neurons of interest can be marked for further quantitative comparisons. Next, level one to three metrics can be compared with paradigm-specific features of the selected neuron(s). For example, in case of hippocampal neurons, place field, trial-by-trial variability of firing patterns, travel direction firing specificity, spike phase precession relative to theta oscillation cycles, and multiple other features can be predefined by the experimenter. During the data mining process, unexpected features and outliers may be noted, instabilities of neurons (‘drifts’) can be recognized, and artifacts identified. Such experimenter-supervised judgments are also essential for evaluating the quality of quantified data processing and estimating potential single neuron-unique features that might drive population statistics.

**Figure 4.**
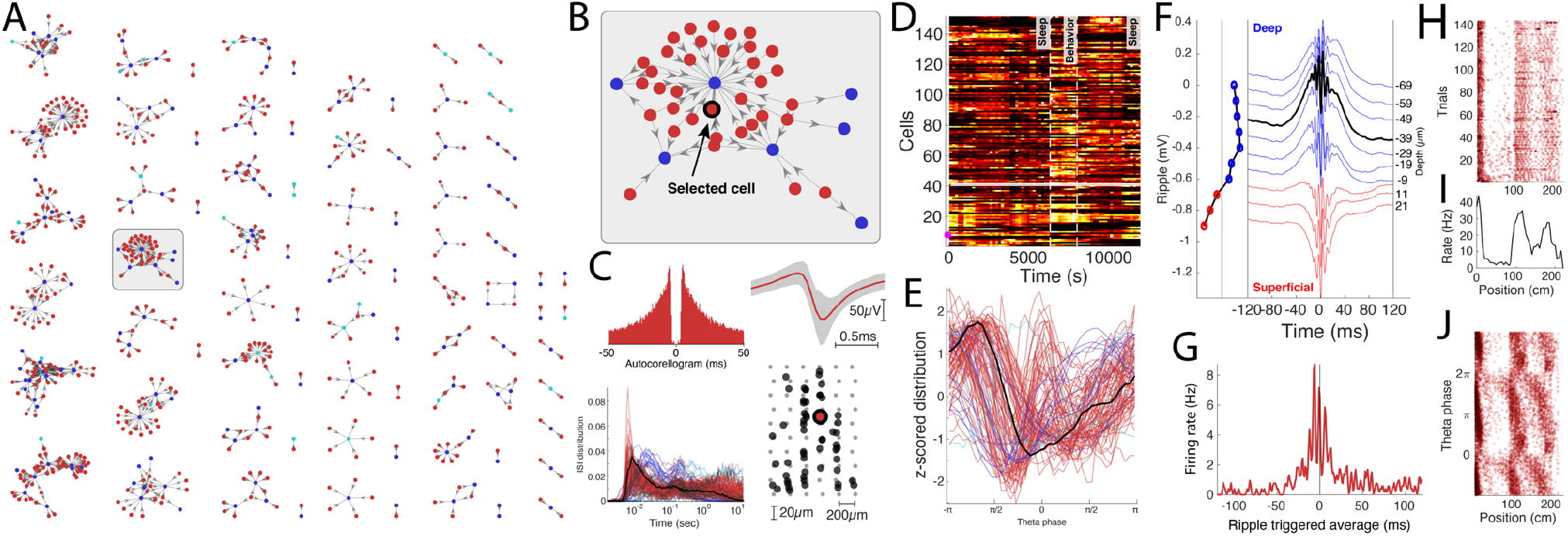
Data exploration example. **A.** Connectivity graph with monosynaptic modules found across multiple datasets. Neurons are color-coded by their putative cell types (pyramidal cells in red, narrow interneurons in blue and wide interneurons in cyan). **B.** Highlighted monosynaptic module with single pyramidal cell highlighted (arrow). **C.** First level metrics: Auto-correlogram, average waveform (top row; gray area signifies the noise level of the waveforms), ISI distributions, with the selected neuron in black, and the physical location of the neurons relative to the multi-shank silicon probe. **D.** Firing rate across time for the population, each neuron is normalized to its peak rate. The session consists of three behavioral epochs: pre-behavior sleep, behavior (track running), and post-behavior sleep (boundaries shown with dashed lines). **E.** Theta phase distribution for all neurons recorded in the same session (red, pyramidal cells; blue, interneurons) during locomotion with the selected neuron highlighted (black line). **F.** Average ripple waveform for the electrode sites on a single shank. The site of the selected neuron is highlighted (dashed black line). The polarity of the average sharp wave is used to determine the position of the neuron relative to the pyramidal layer in CA1. **G.** Ripple wave-triggered PSTH for the selected neuron aligned to the ripple peak. **H.** Trial-wise raster for the selected neuron in a maze. **I.** The average firing rate of the neuron across trials. **J.** Spike raster showing the theta phase relationship to the spatial location of the animal. All of the place fields show phase precession.

Several benchmarks were performed to characterize the performance of CellExplorer under various conditions, benchmarking single cell plots, and computation of various features. The results are summarized in Supplementary Fig. 5.

### Value of large inter-laboratory datasets

While progress in discovery science often depends on the investigator-unique approach to novel insights, standardization of data processing and screening is essential in fields where ‘big data’ generation is achieved through collaborative efforts. This applies to the current effort to quantitatively relate physiology-based and genetically classified cell types (Klausberger and Somogyi, 2008; McBain and Fisahn, 2001; Rudy et al., 2011). In each experiment, typically only one or a limited number of neuron types can be identified. Yet, combining datasets from numerous experiments and different laboratories can generate physiological metrics, grounded by optogenetics and other ‘ground truth’ data.

Fig. 5 illustrates the feasibility and utility of this approach. Level one to three metrics of neurons recorded from the same brain region and layer can be combined from multiple experiments and laboratories and contrasted to the data quality of units recorded in a single session. An ever-growing data set allows for more reliable modality separation and characterization of neuron types. For example, the initial divisions of neurons into putative pyramidal cells, narrow and wide interneurons can be further refined by quantifying monosynaptic connections, increasing confidence of pyramidal cell–interneuron separation as well as identifying subsets of the unclassified group as interneurons (Fig. 5a) (Mizuseki et al., 2011; Petersen and Buzsáki, 2020; Peyrache et al., 2015; Stark et al., 2013). Combining extracellular spiking with intracellular recordings can further help determine the excitatory or inhibitory identity of neurons (Radosevic et al., 2019).

**Figure 5.**
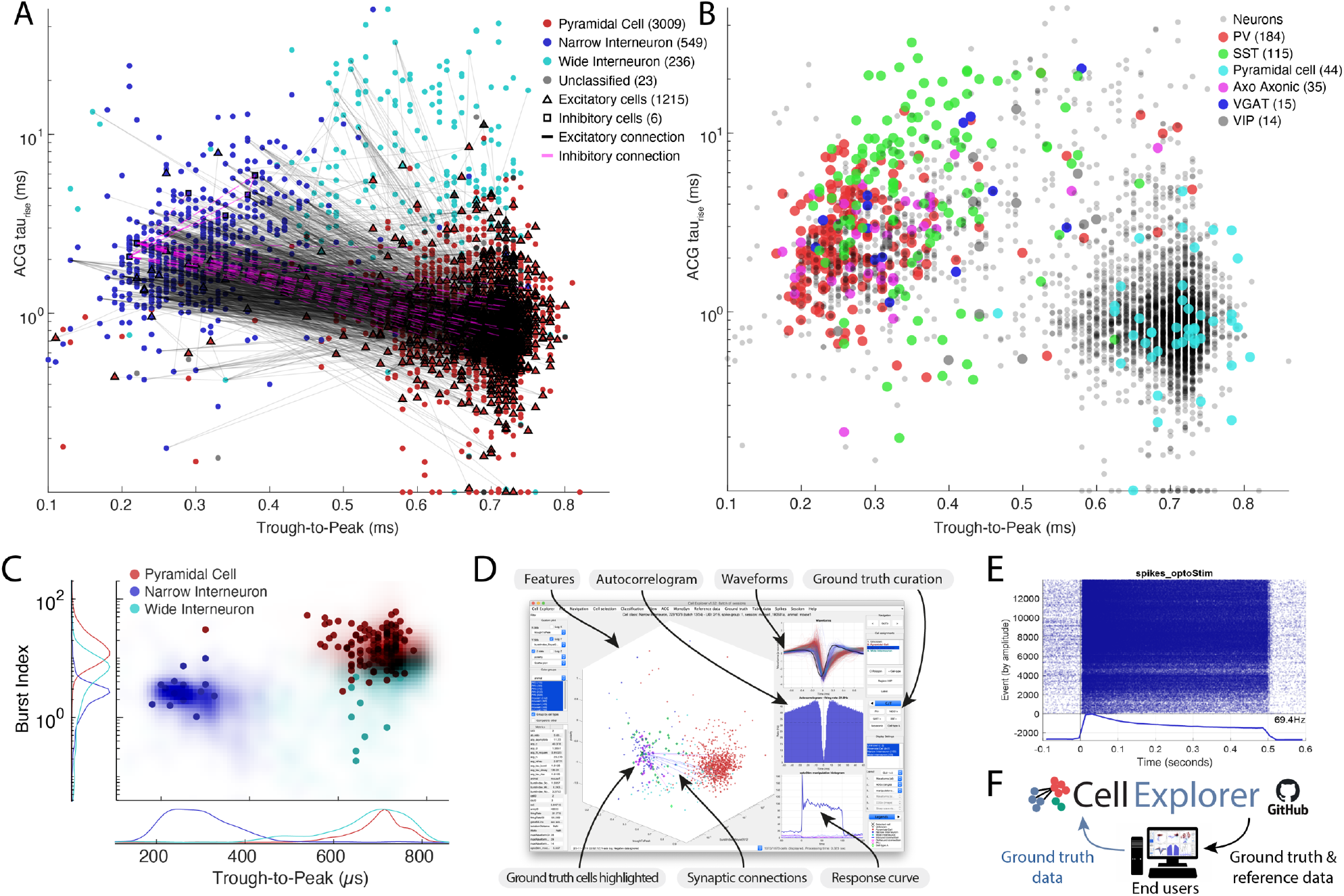
Ground truth data, reference data, and opto-tagging. **A.** Distribution of putative cell types (3811 cells), including their projections determined via spike-transmissions CCG curves (Petersen and Buzsáki, 2020; Petersen et al., 2020). **B.** 407 ground truth neurons spanning PV (184), SST (115), Pyramidal cells (44), Axo-axonic (35), VGAT (15) and VIP cells (14) projected on the same population of neurons as in A. Data from ref. Petersen et al. (2020) and Allen Brain Observatory Neuropixels database (https://portal.brain-map.org/explore/circuits/visual-coding-neuropixels; Siegle et al., 2019). **C.** Single session (dots) compared with data from 30 reference sessions (shaded zones). D. Opto-tagged data can be processed and curated directly in CellExplorer. **E.** Example of a PSTH of a PV-expressing neuron to 500 ms square light pulses. Raster plot and average responses to the light pulses visualized in CellExplorer. **F.** The CellExplorer framework allows for sharing ground truth and reference data directly with the end-user. End users can upload their ground truth data to the CellExplorer GitHub repository for communal sharing (see the opto-tagging tutorial at CellExplorer.org).

Single neurons identified by opto-tagging or other direct means (Ciocchi et al., 2015; Klausberger and Somogyi, 2008; Royer et al., 2012; Senzai et al., 2019; Stark et al., 2012; Zhang et al., 2013; Roux and Buzsáki, 2015) can be used to link level one to three features of initially classified neurons to genetically defined neuron types (Fig. 5b). For example, expressing a light-gated ion channel under the control of a specific promotor makes it possible to evoke extracellularly detectable action potentials in GABA transporter (VGAT) expressing inhibitory interneurons, and even in interneuron sub-types, including parvalbumin (PV), somatostatin (SST), vasoactive intestinal peptide (VIP), vesicular, chandelier cells (axo-axonic cells), as well as in pyramidal cells (Fig. 5b). Having access to these ground truth labels may offer further clues for a separation based on physiological criteria. An expected outcome of the growing number of datasets containing ground truth-verified neurons, is trained models for classifying diverse neuron types based on physiological metrics alone. This is especially important for recordings in model organisms for which genetic manipulations are less tractable than in mice. Opto-tagged neurons can be explored in CellExplorer, e.g. by displaying the average PSTH (Figure 5D) as well as trial-wise raster plots (Figure 5E). Further manual curation can be done while accessing the neuron’s other characteristics including waveforms, firing rates and connectivity. Communal contribution of ground truth data to CellExplorer is possible through the public GitHub repository (Figure 5F; visit CellExplorer.org for tutorials and further details).

CellExplorer uses and shares data through our lab databank (buzsakilab.com/wp/database) (Petersen et al., 2020). Through our database and CellExplorer, we currently share more than 79.000 processed neurons publicly. The datasets are all from awake rodents, from freely behaving rats and mice in our laboratory, as well as neurons from two large datasets from head-fixed mice: the Allen Institute Visual Cortex dataset (Siegle et al., 2019) and the UCL dataset spanning many brain regions (Steinmetz et al., 2019). All these datasets can be downloaded directly in CellExplorer and used as reference data or explored directly (Fig. 6). The infrastructure is designed towards continually growing the public datasets and ground truth data with the goal of building ever larger data banks for discovery science, cross-laboratory interactions, and reproducibility control.

**Figure 6.**
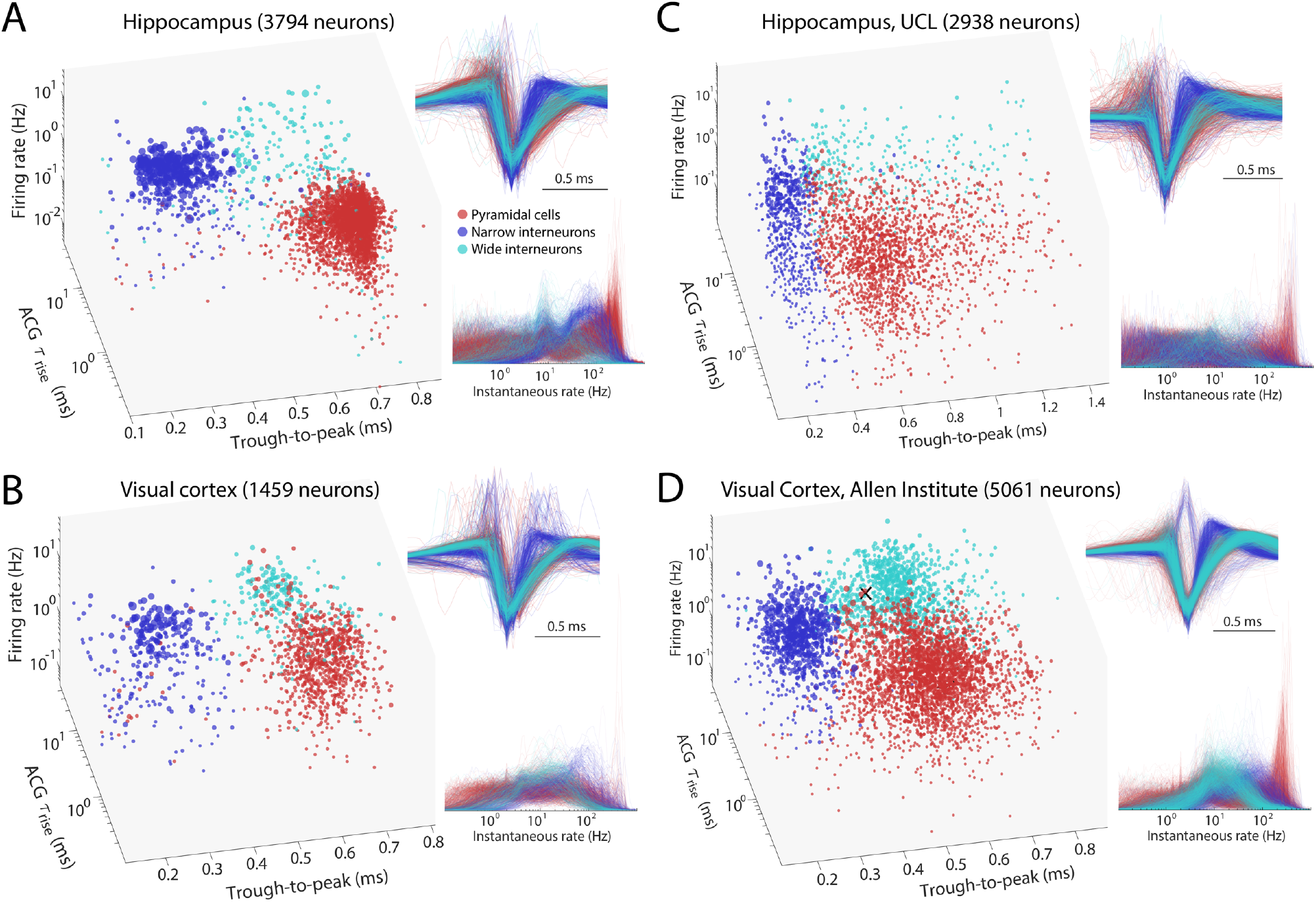
Comparison of initial neuron classification by CellExplorer on large scale datasets from three different laboratories. **A.** Data from hippocampus (Petersen and Buzsáki, 2020). **B.** Data from visual cortex (Senzai et al., 2019). **C.** Hippocampal and visual neurons selected from the UCL dataset (Steinmetz et al., 2019). **D.** Visual cortex cells from the Allen Institute (Siegle et al., 2019). Right panels across A-D: Z-scored waveforms across all neurons (top) and distribution of instantaneous rates (1/interspike intervals) across all neurons. A and B are based on long homecage (sleep) data (several hours), while C and D data are from short (~ 30 min) sessions in head-fixed, task-performing mice. See also Suppl. Fig. 6. Red, pyramidal cells; Blue, narrow waveform interneurons; Cyan, wide waveform interneurons.

Using large numbers of shared datasets, one can begin to compare brain regions, different electrode types, and other features efficiently using t-SNE plots. Such representations can highlight inconsistencies and differences across recording sessions, identify important regional and layer-specific differences, and alert for interspecies characteristics (Supplementary Fig. 6).

### Demonstration of the inter-laboratory applicability of CellExplorer

To demonstrate the applicability of CellExplorer, we processed datasets from the CA1 region of the hippocampus (Figure 6A) and visual cortex (Figure 6B) in freely moving mice (Senzai et al., 2019; Petersen and Buzsáki, 2020; Petersen et al., 2020), and comparable data from the two largest public dataset collections from University College London (Fig 6C;Steinmetz et al., 2019) and the Allen Institute (Figure 6D; Siegle et al., 2019). Processing data collected in different laboratories and investigators by the same program(s) will allow investigators to standardize protocols for levels one to three metrics and, therefore, higher reliability of interlaboratory experiments between neuronal firing patterns and their behavioral, cognitive correlates. For example, baseline data collected for levels one to three classification, such as bursts, firing rates, coefficient of variation of interspike intervals, may differ whether the recordings are made during active behavior, immobility or sleep.

## DISCUSSION

We have developed CellExplorer, a transparent, open-source, MATLAB-based resource for characterizing single neurons and neuron types based on their electrophysiological features. The CellExplorer platform enables visualization and analysis for users without the need to write code. Its modular format allows for fast and flexible comparisons of a large set of preprocessed physiological characteristics of single neurons and their interactions with other neurons, as well as their correlation with experimental variables. The code is publicly available on GitHub for users to download and to use the same standardized processing module on their local computers (Windows, OS X, and Linux).

CellExplorer offers step-by-step online tutorials for first-time users. It is linked to the Allen Institute reference atlas (Chon et al., 2019; Wang et al., 2020; https://atlas.brain-map.org/) and can be expanded to include other online resources that provide annotated data on putative neuron types. The open-source design of CellExplorer permits the optimization of current standardized components tested against the ever-growing community-contributed database and will accelerate efforts to link genetically identified neuron types with their physiological properties in the intact brain.

### Multiple-level characterization and classification of single neurons

To correctly interpret neuron firing-behavior/cognition relationships, numerous controls are needed to rule out or reduce the potential contribution of spurious variables. The Processing Module generates a battery of useful metrics for this. In addition to the first-level description of the biophysical and physiological characteristics of single neurons, it can compute brain state-dependent firing rates, interspike interval variation, and relationships between single neurons and spiking activity of the population and LFP (second level). When behavioral data is also available, it can describe the relationship between single neuron firing patterns and routine behavioral parameters, such as immobility, walking, respiration, and pupil diameter (third level). The third-level metrics can help avoid inappropriately attributing spiking activity to high-level phenomena, such as learning, perception or decision making, that are often linked to overt movement and autonomic changes. Because these three-level metrics are independent of particular experimental paradigms, they can be used as benchmarks for assessing consistencies across experiments performed by different investigators in the same laboratory or across laboratories (Figure 6). Concatenating datasets obtained from the same brain regions and layers will create a continuously growing data bank. In turn, these data-rich sets make it possible to identify and quantify reliable boundaries among putative clusters and suggest inclusion and exclusion of parameters for a more refined separation of putative neuronal classes. Sets from different brain regions can be readily compared in order to identify salient differences.

Although several statistical tests are available in CellExplorer, it is not meant to substitute rigorous quantification. Instead, it is designed as a tool for facilitating visualization, interpretation and discovery. It is a complementary approach to dimensionality reduction and population analysis methods. Because assemblies of neurons consist of highly unequal partners (Buzsáki and Mizuseki, 2014), knowledge about the neuron-specific contribution to population measures is critical in many situations (Nicolelis and Lebedev, 2009). Such inequality may stem from unknowingly lumping neurons of different classes together into a single type and because even members of the same type belong to broad and skewed distribution and may contribute to different aspects of the experiment (Grosmark and Buzsáki, 2016).

Two key features make the CellExplorer platform highly efficient: flexibility and speed. Flexibility is provided by the numerous parameters as outputs of the Processing Module. High speed is achieved by using pre-computed metrics and limiting the computation time in the user interface. A caveat is that online alteration of the plots and metrics is not allowed. Yet, such changes can be performed by working with the raw spike data. Multiple features of single neurons, displayed on the same screen, can be compared. Moving from one neuron to the next and its multiple displayed features requires only a mouse click. These multiple features can be compared by superimposing the same features of two or more neurons, all neurons in a session, or the entire database. This allows unexpected features to be noted, or obvious artifacts to be recognized and deleted. When unusual sets are discovered in any display, all other features of the same set can be rapidly compared and contrasted to other sets. Neuron clouds can be selected by drawing polygons around them and regrouped in any arbitrary configuration. Inspection of datasets containing even several thousand neurons (Siegle et al., 2019; Steinmetz et al., 2019) is realistic because minimal computing time is required in the graphical interface and because in most conditions only small subsets need individual inspection and quality control.

Various classification schemes have been developed to assign extracellular spikes to putative pyramidal cells, interneurons, and their putative subtypes, based on a variety of physiological criteria. These include waveform features, firing rate statistics in different brain states, embeddedness in various population activities, firing patterns characterized by their autocorrelograms, and putative monosynaptic connections to other neurons (Barthó et al., 2004; Csicsvari et al., 1999; Fujisawa et al., 2008; Mizuseki et al., 2009; Okun et al., 2015; Sirota et al., 2008). Increasingly larger datasets will likely improve such physiology-based classification. Yet, the ‘ground truth’ for these classifying methods is largely missing. Optogenetic tagging (Boyden et al., 2005) offers such grounding by connecting putative subtypes based on physiologically distinct features to their molecular identities. Because in a single animal only one or few neuron types can be tagged optogenetically or identified by other direct methods (Fosque et al., 2015; Klausberger and Somogyi, 2008), refinement of a library of physiological parameters should be conducted iteratively, so that in subsequent experiments the various neuron types can be recognized reliably by using solely physiological criteria (English et al., 2017; Royer et al., 2012; Senzai and Buzsáki, 2017, 2017; Roux and Buzsáki, 2015). In turn, knowledge about the molecular identity of the different neuronal components of a circuit can considerably improve the interpretation of correlational observations provided by large-scale extracellular recordings.

### Data sharing

Above, we described one of the many possible examples that can benefit from large databases. Currently, tens to hundreds of thousands of pre-processed neurons exist across laboratories in different brain regions, which can be identically streamlined by the Processing Module, and displayed and compared in the same coordinate system. We welcome shared datasets from other research groups for enhancement and comparison with our publicly-accessible database (buzsakilab.com/wp/database; Allen Institute; UCL; Petersen et al., 2020; Siegle et al., 2019; Steinmetz et al., 2019). The single prerequisite for quantitative comparison of data across laboratories is to make wideband data available (≤ 3 Hz to ≥ 8 kHz; ideally ≥ 20 kHz sampling rate) so that all data are processed the same way.

Through our web resource (Petersen et al., 2020), we host > 1,000 publicly shared datasets of long (4 to 24 hrs), large-scale recordings of single units from multiple brain structures, including the hippocampus, entorhinal, prefrontal, somatosensory, and visual cortices, thalamus, amygdala, and septum (buzsakilab.com/wp/database). The database also includes the data from Allen Institute and UCL. Long-recordings have the advantages of defining brain state-dependent characteristics of neurons, such as their firing rates and patterns during waking and sleep, unmasking the ‘hidden’ or a relatively silent majority of neurons (Mizuseki and Buzsáki, 2013; Shoham et al., 2006) and discovering their connectivity patterns (English et al., 2017). These data already provide benchmarks assessing the reliability of initial neuron classification into the broad groups of pyramidal cells and interneurons, many of which are identified physiologically by their monosynaptic connections. They also offer normative data about spikes features, firing rates, and spike dynamics. These features can serve as benchmarks for comparison with data collected in any other laboratory.

### Development and availability

Development takes place in a public code repository at github.com/petersenpeter/CellExplorer. All examples in this article have been calculated with the pipeline and plotted with CellExplorer. Extensive documentation, including installation instructions, tutorials, description of all metrics and their calculations, is hosted at CellExplorer.org. CellExplorer is available for MATLAB 2017B and forward, and for the operating systems Windows, OS X, and Linux. More information can be found at CellExplorer.org. All data presented is available from https://buzsakilab.com/wp/database/ (Petersen et al., 2020).

## ACKNOWLEDGEMENTS

We thank Sam McKenzie, Mihály Vöröslakos, Michelle Hernandez, Thomas Hainmueller, for helping with documentation or implementation of CellExplorer code. Further, we thank Sam McKenzie, Daniel English, Roman Huszar, and Yuta Senzai for providing ground truth datasets. Finally, thanks go to Thomas Hainmueller, Manuel Valero, Antonio Hernandez-Ruiz, Viktor Varga, and Omid Yaghmazadeh for comments on the manuscript. Joshua H. Siegle thanks the Allen Institute founder, Paul G. Allen, for his vision, encouragement, and support. Supported by U19 NS107616, U19 NS104590, R01 MH122391, and The Lundbeck Foundation.

## SUPPLEMENTARY SECTION

**Supplementary figure 1:**
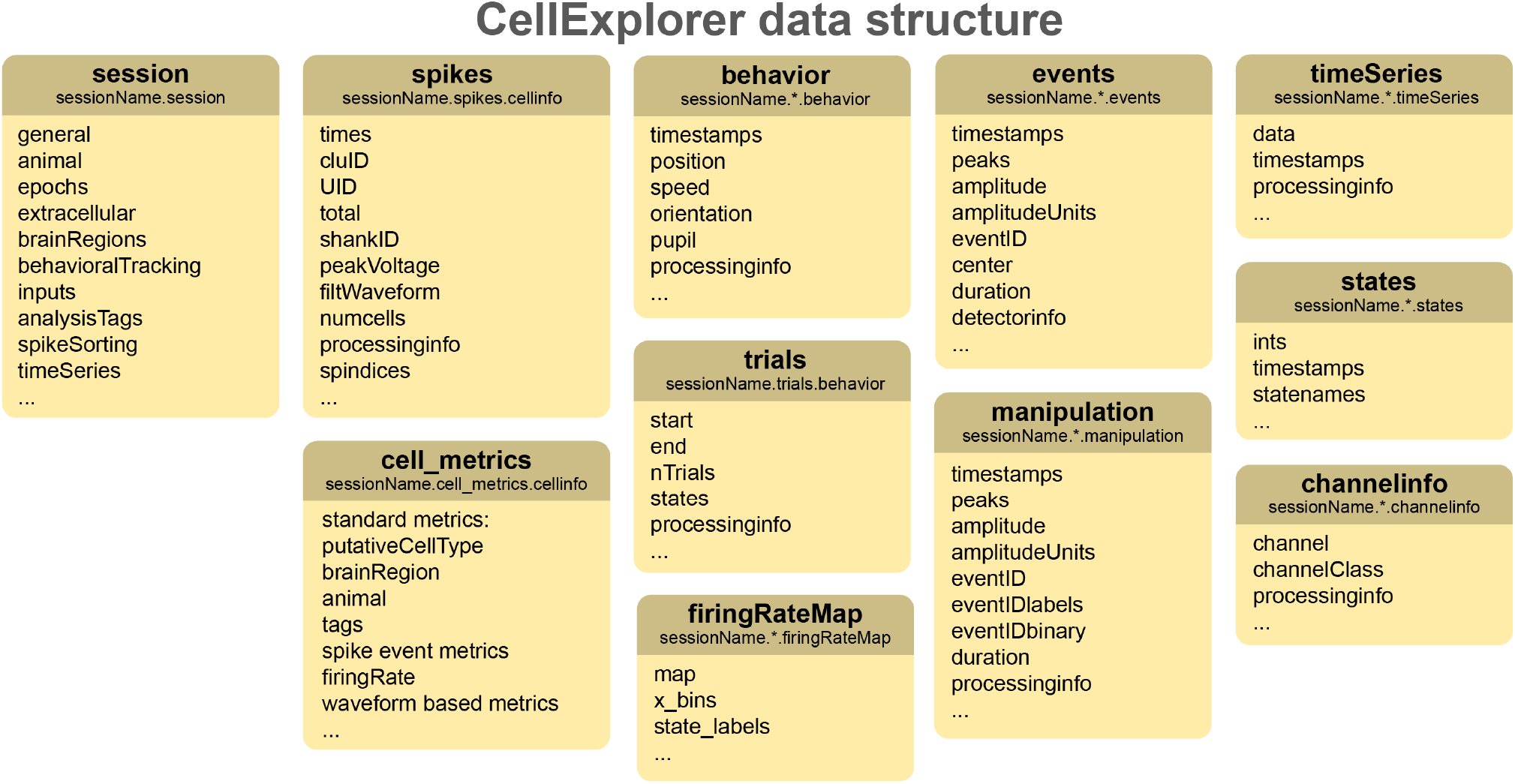
Datatypes. The data structure. A detailed description is available online at CellExplorer.org/datastructure/data-structure-and-format. session, spikes, cell_metrics, trials are defined data types, while behavior, firingRateMap, events, manipulation, timeseries, states, channelinfo are data containers.

**Supplementary Figure 2.**
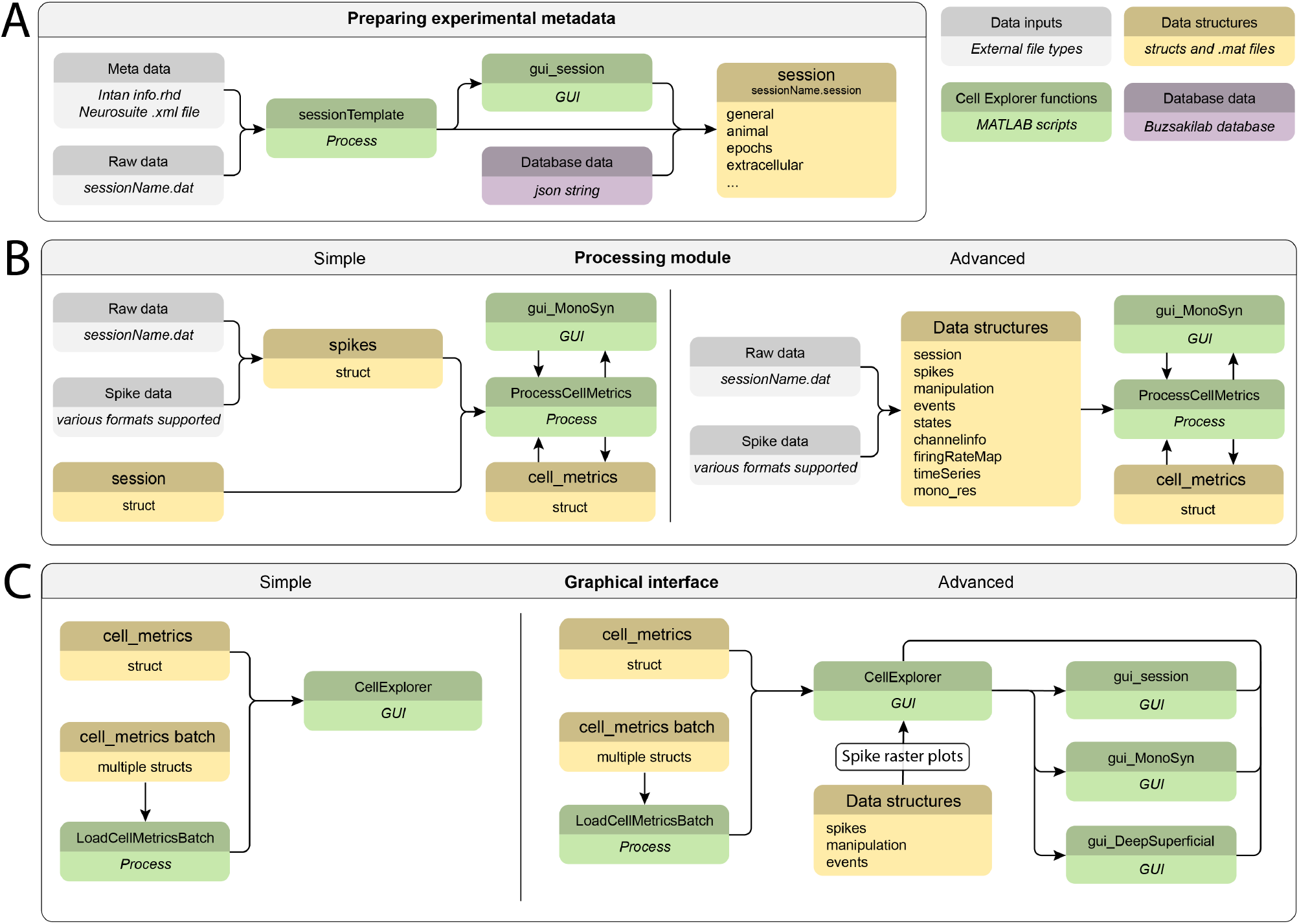
Flow charts. **A)** Generating the metadata structure for a recording session. **B**) Running the processing pipeline. **C**) Running the CellExplorer module for manual curation and exploration. CellExplorer data structures are shown in yellow, MATLAB functions in green, and the input data in grey. Input from the Buzsáki lab database is shown in purple (buzsakilab.com/wp/database).

**Supplementary Figure 3.**
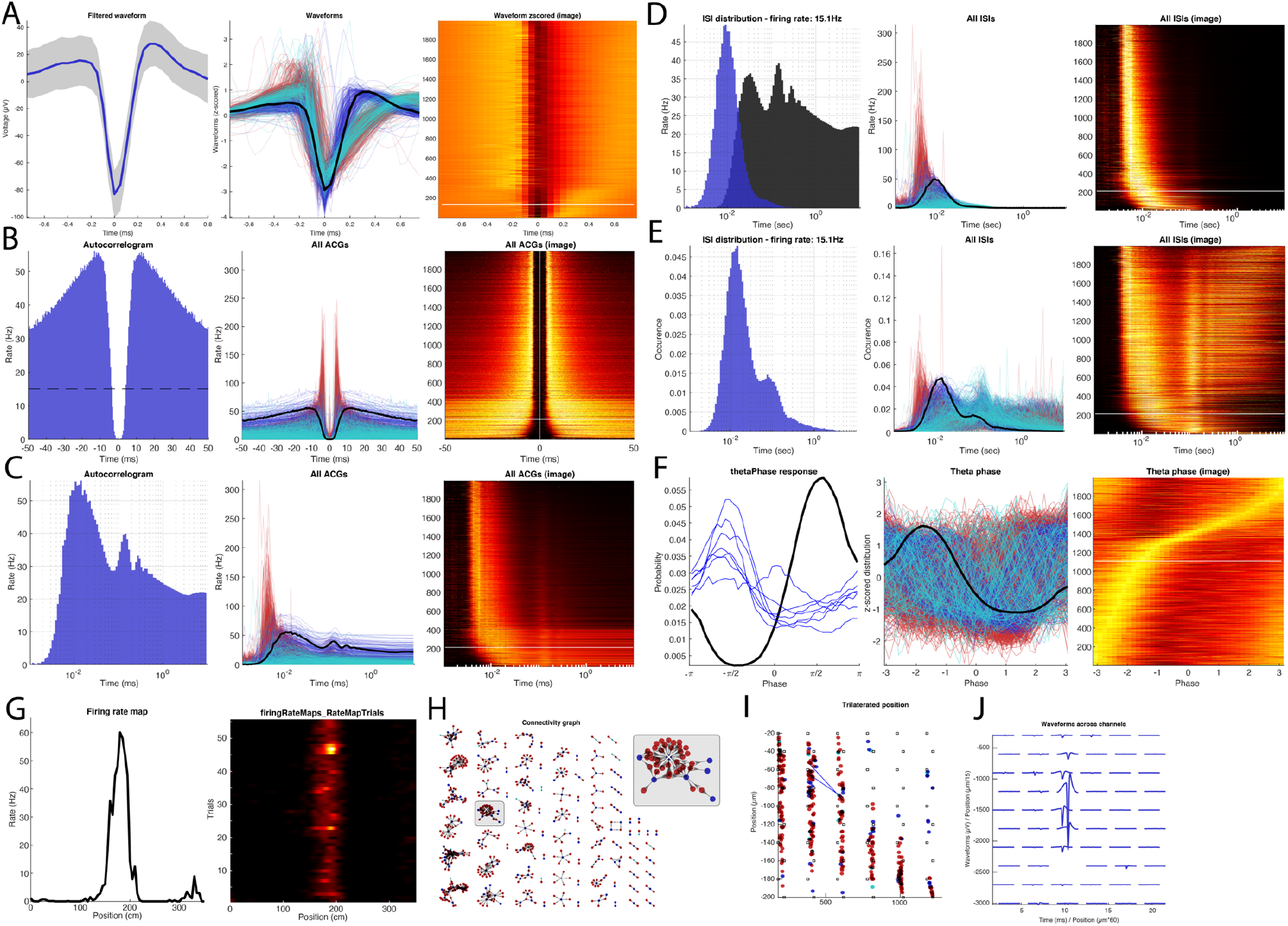
The various single-cell plots. Most single cell data-visualizer have three representations: single neurons (with neuronal connections highlighted for a subset of the plots), all neurons (absolute or normalized representations), and an image representation (normalized data, with selected cell highlighted by a white line). **A.** Waveform representations: waveform of a selected single neuron, waveforms of all neurons (z-scored), and their image representation. The white line in the image representation corresponds to the selected neuron. **B.** Autocorrelograms (ACGs) for the single neuron, ACGs for all neurons and their image representation. **C.** ACGs on a log scale (single, all, image). **D, E.** Interspike interval distributions (ISIs) on a log scale (single, all image) for two different normalizations (**D,** rate (Hz); **E,** occurrence). **F.** Theta phase spike histogram for the single interneuron (black line) and those of pyramidal neurons monosynaptically connected to the interneurons (blue lines; left) and all neurons in the same session (middle and right panels). **G.** Firing rate map for a pyramidal cell. Session average (left) and trial-wise heatmap. **H.** Connectivity graph showing all monosynaptic modules in the dataset. A module is highlighted and enhanced (top right). **I.** Physical location of neurons recorded in the same animal using trilateration. Eight-shank silicon probe recording (8 sites on each shank). Red, pyramidal cells. Blue, interneurons. Monosynaptic connections between two pyramidal cells and a target interneuron are also shown (blue lines) **J.** Average waveform across channels of the single interneurons shown in most panels. A-F, H-J: a narrow interneuron, G: Spatial firing rate of a pyramidal cell on a linear track.

**Supplementary Figure 4.**
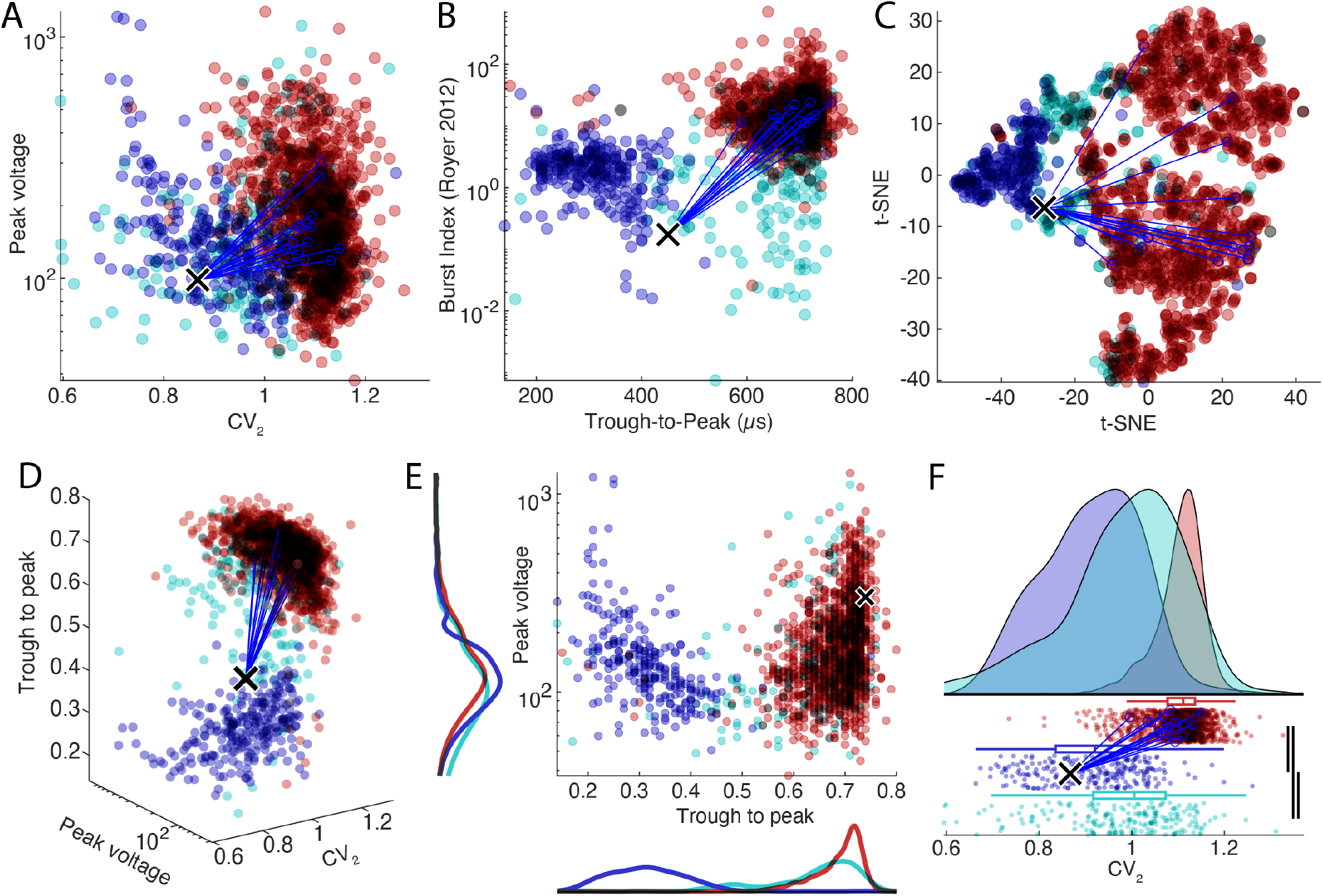
Population data plots. Top row: The three standard representations: custom plot (**A**), classic representation (**B**), and t-SNE plot (**C**). **Bottom row:** The custom plot has 3 further data representations: a 3-dimensional plot with custom marker size (**D**), 2D plot with marginal histograms (**E**), and one-dimensional raincloud plots (**F**), combining 1D scattered neurons with error bars histogram and KS significance test (line thickness represent significance levels). Color-coded according to cell types: pyramidal cell (red), narrow interneuron (blue), wide interneuron (cyan).

**Supplementary figure 5.**
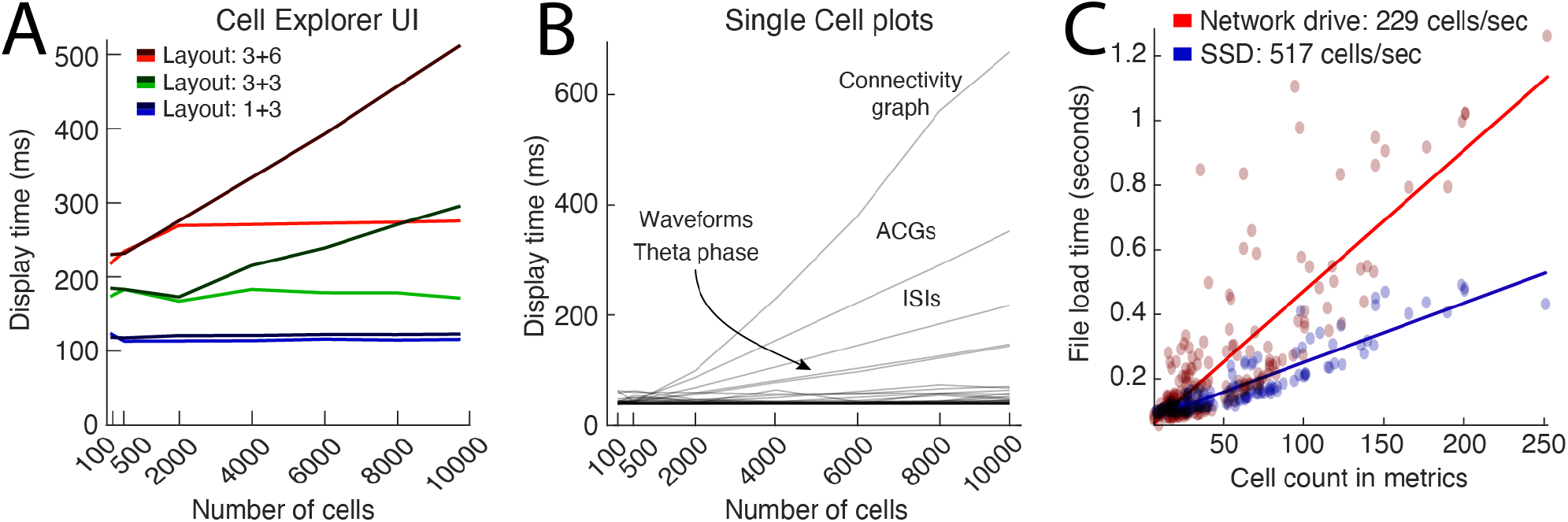
Benchmarks of the CellExplorer user interface (UI). **A.** UI display times when switching between units for the three layouts shown in figure 3B (approximately 110 ms for layout 1+3; blue lines. 180 ms for layout 3+3; green lines) and 290 ms (layout 3+6; in red), respectively. Dark gradient colored lines (dark red, green, and blue) indicate where there were no limits on the number of traces plotted for single-cell plots, and the light gradient lines show screen update times with a maximum of 2000 random traces. **B.** Display times for single-cell plots, quantified by the number of cells displayed. The plots contributing most to an increased display time are the plots with trace representations for each cell (ACGs, ISIs, waveforms, ISIs, theta phase) and the connectivity graph. By default, a maximum of 2000 traces is drawn capping the processing time below ~80 ms for all plots except the connectivity graph. **C.** Benchmarking of cell metrics file readings. On average, 230 cells can be loaded per second quantified across 180 sessions with various cell count (red dots and linear fit in red). By storing the data on a local SSD, the loading time could be decreased and attain cell loading above 500 cells per second.

**Supplementary figure 6.**
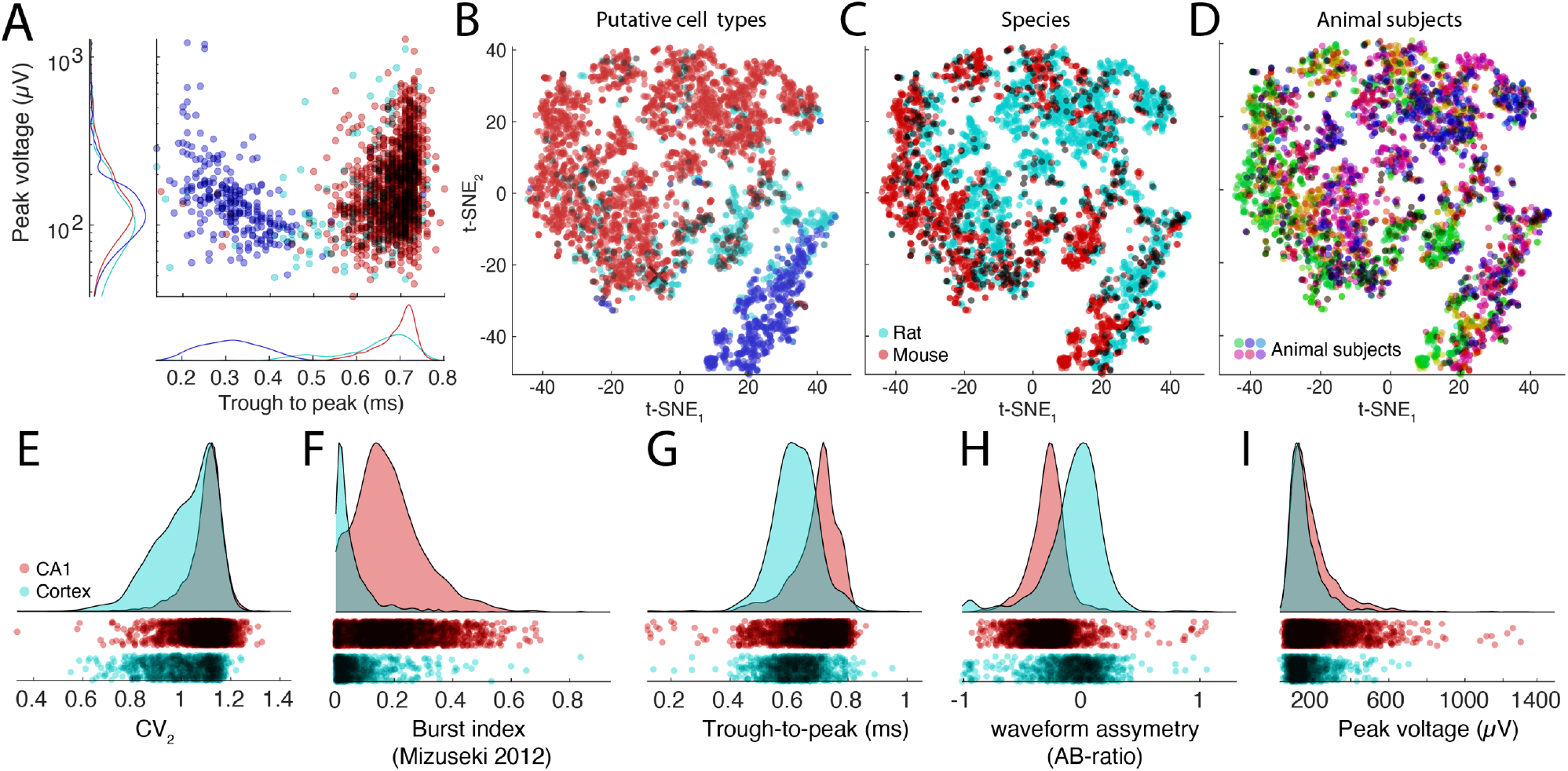
Exploration and comparison of metrics and cells across, species, subjects and brain regions. **A.** Distributions of spike amplitudes and waveform width (quantified by the trough to peak metrics) for the three groups from multiple CA1 datasets. Note inverse relationship between spike amplitude and waveform for putative interneurons. **B-D.** t-SNE representations of putative cell types (B), species (C, rat, and mouse in magenta and red, respectively) and subjects (D, colors scaled across subjects) for hippocampal neurons. **E-I:** Comparison of spike features of neurons recorded from CA1 pyramidal cells and visual cortex pyramidal cells. Significant differences are observed across several basic metrics, including CV_2_ (E), burst index (F), trough-to-peak (G), waveform asymmetry (H), and waveform peak voltage (I).

**Supplementary figure 7.**
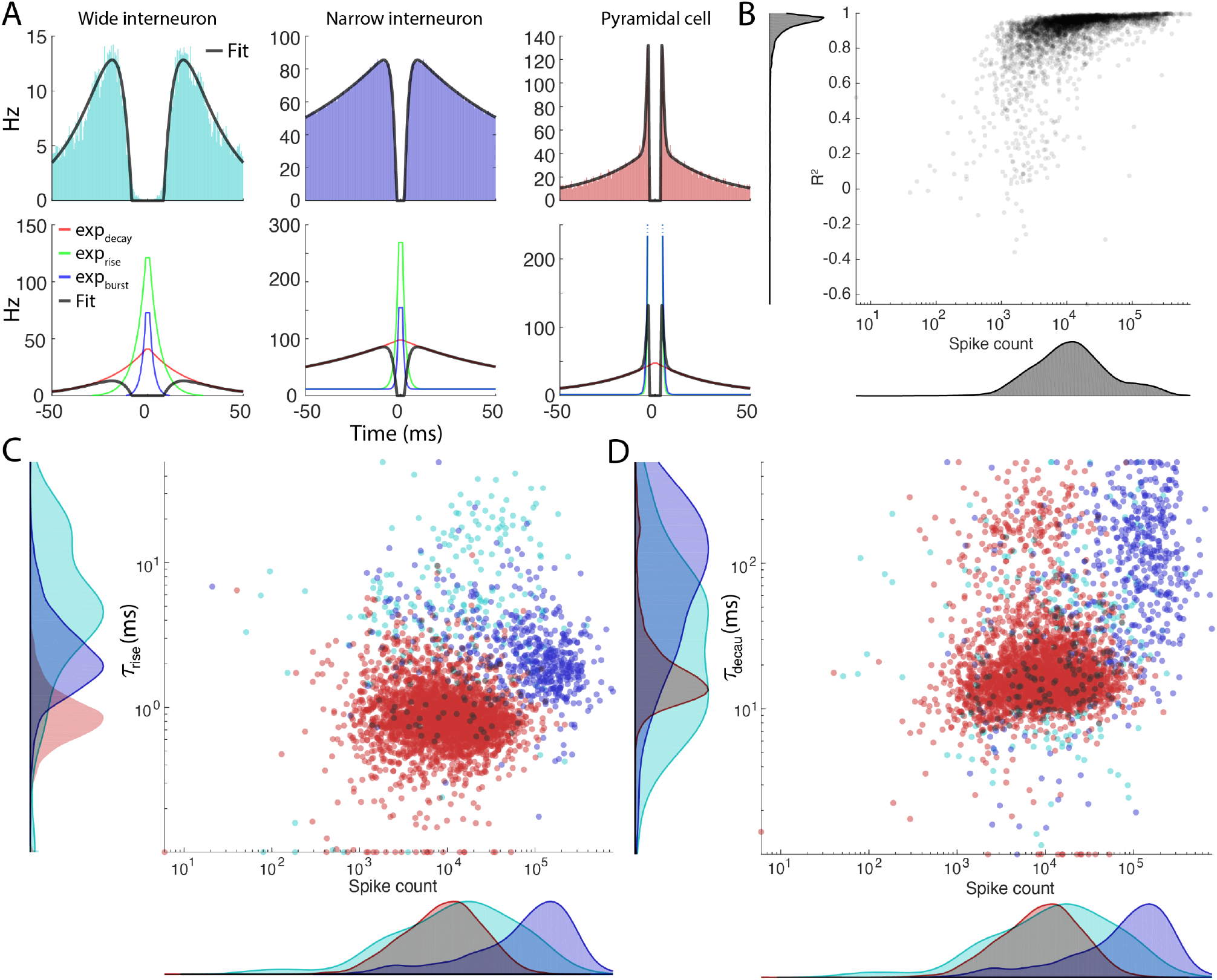
ACG fits. **A.** Three examples of typical autocorrelograms for a wide interneuron (left column) narrow interneuron (middle column) and a pyramidal cell (right column). The exponential components are plotted in the lower row. **B.** The R^2^ values for each fit across the 4000 pyramidal cells plotted against the number of spikes. **C-D *τ***_rise_ (C) and ***τ***_decay_ (D) values plotted against the spike count. Color coded by putative cell type.

**Supplementary Table 1.** Primary MATLAB functions of the CellExplorer framework.

| Functions | Description |
| --- | --- |
| sessionTemplate | A template script which automatically extracts and imports relevant metadata |
| gui_session | A graphical user interface (GUI) for manual inspection and entry of metadata |
| ProcessCellMetrics | The processing module |
| CellExplorer | The main graphical interface of CellExplorer |
| preferences_CellExplorer | Preferences for the graphical interface |
| preferences_ProcessCellMetrics | Preferences for the processing module |
| gui_MonoSyn | GUI for manual curation of monosynaptic connections |
| gui_DeepSuperficial | GUI for manual curation of the depth assignment of neurons based on depth-related changes of sharp-wave-ripples (Mizuseki et al., 2011) |
| loadCellMetricsBatch | Batch loading script for combining cell_metrics structs across sessions |
| loadCellMetrics | Script for loading cell_metrics with built-in common text filters (putative cell type, brain region, synaptic effect, label, animal, tags, groups, etc.) |

**Supplementary Table 2:**
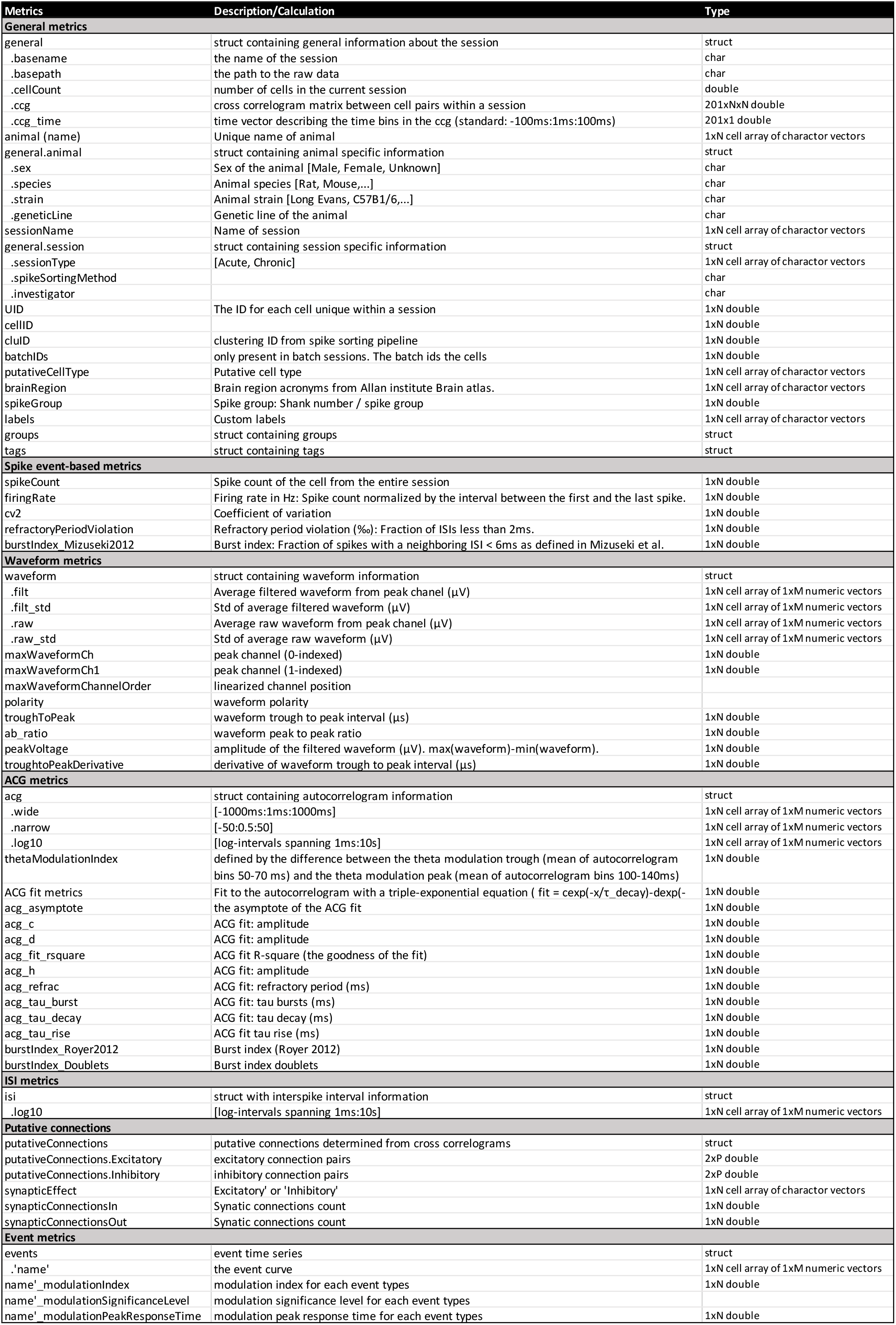
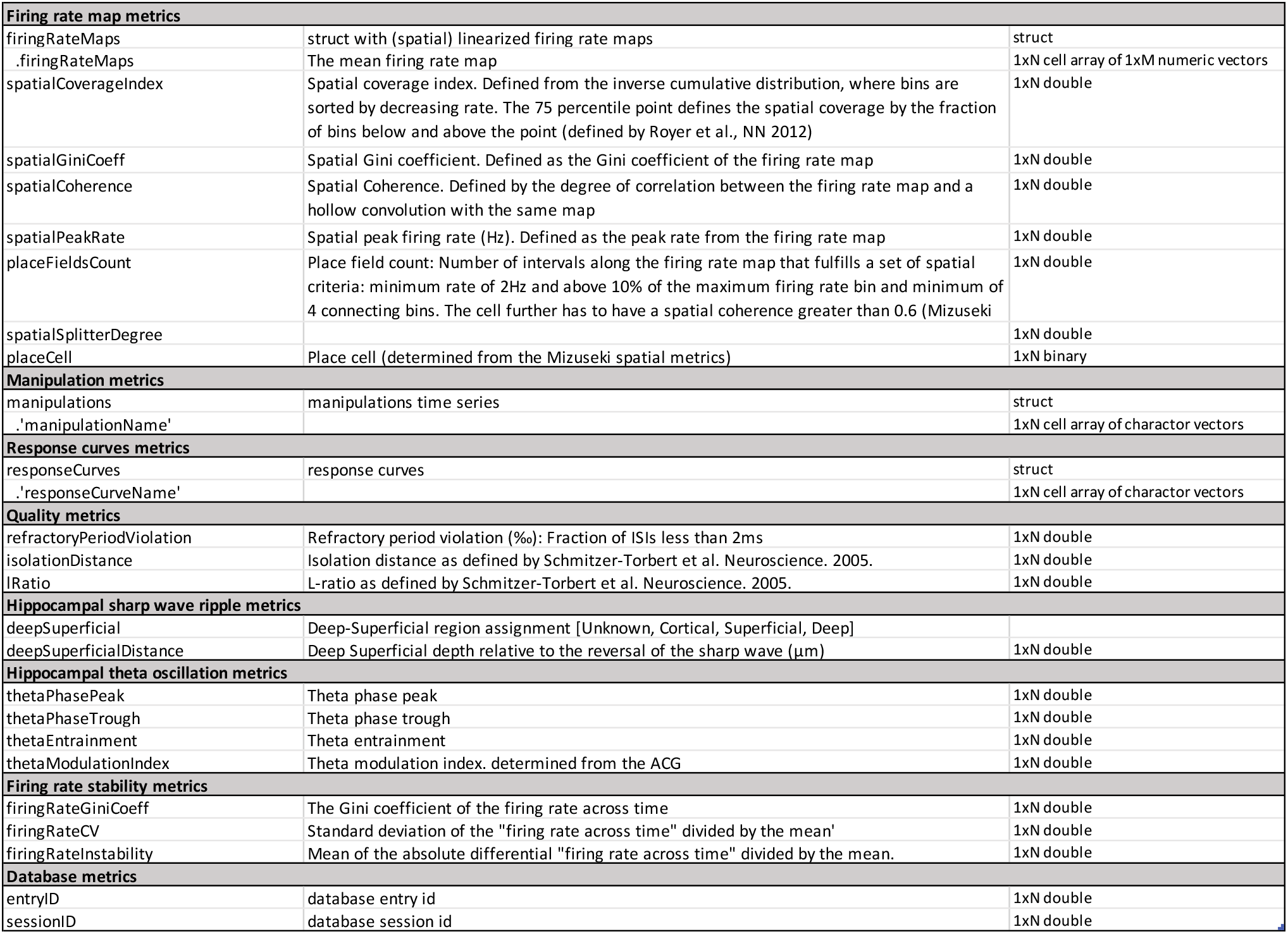
Cell metrics. An incomplete list of the standard cell metrics. The full list is available online at CellExplorer.org/datastructure/standard-cell-metrics

**Supplementary video 1.**
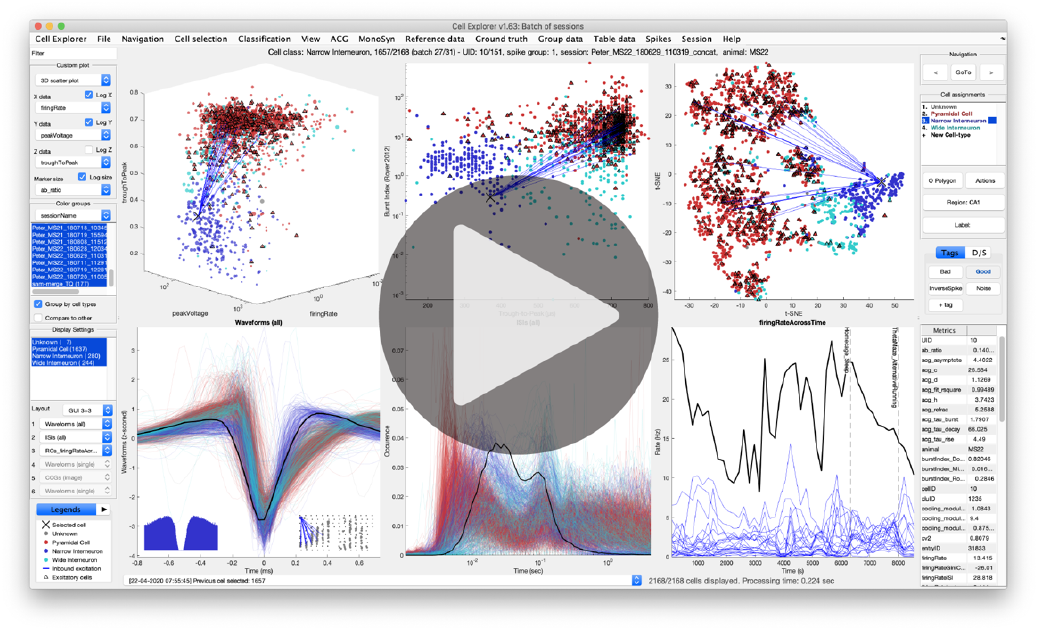
The supplementary video is available on <u>YouTube</u> and at CellExplorer.org.

## TUTORIALS

A general tutorial on the full pipeline is available below. There are many more detailed tutorials online, covering: the generation of the metadata struct, the manual curation process, generating spike raster plots, connections, performing opto-tagging, using ground truth data, export figure, and many other topics.

Tutorials are available online at CellExplorer.org/tutorials/tutorials

## General tutorial

This tutorial shows you the full processing pipeline, from generating the necessary session metadata using the template, running the processing pipeline, opening multiple sessions for manual curation in CellExplorer, and finally using the cell_metrics for filtering cells, by two different criteria. The tutorial is also available as a Matlab script: (`tutorials/CellExplorer_Tutorial.m`).

1. Define the basepath of the dataset to process. The dataset should consist of a `basename.dat` (a binary raw data file), a `basename.xml` (recommended; not required) and spike sorted data.

~~~
basepath = ‘/your/data/path/basename/’;
cd(basepath)
~~~
2. Generate session metadata struct using the template function and display the metadata in a GUI

~~~
session = sessionTemplate(basepath, ‘showGUI’, true);
~~~ In the GUI you can put in relevant metadata. Please pay attention to the general, extracellular, and spike sorting tabs and verify all metadata.
3. Run the cell metrics pipeline `ProcessCellMetrics` using the session struct as input

~~~
cell_metrics = ProcessCellMetrics(‘session’, session);
~~~
4. Visualize the cell metrics in CellExplorer

~~~
cell_metrics = CellExplorer(‘metrics’, cell_metrics);
~~~
5. Now you can repeat step 1-4 on a couple of datasets and load them together in CellExplorer, providing several paths

~~~
basepaths = {‘path/to/session1’,‘path/to/session2’};
cell_metrics = loadCellMetricsBatch(‘basepaths’, basepaths);
cell_metrics = CellExplorer(‘metrics’, cell_metrics);
~~~
6. Curate your cells and save the metrics
7. Finally, to incorporate the cell metrics into your analysis you can use the load function that has filters built-in:

1. Get cells that are assigned as *Interneuron*

~~~
cell_metrics_idxs1 = loadCellMetrics(‘cell_metrics’, cell_metrics, ‘putativeCellType’, {‘Interneuron’});
~~~
2. Get cells that have the groundTruthClassification label *Axoaxonic*

~~~
cell_metrics_idxs2 = loadCellMetrics(‘cell_metrics’, cell_metrics, ‘groundTruthClassification’, {‘Axoaxonic’});
~~~

